# Microbial and organic matter composition jointly drive phosphorus cycling genes and phosphorus availability in Amazonian soils

**DOI:** 10.1101/2025.10.14.682339

**Authors:** Guilherme Lucio Martins, Gabriel Gustavo Tavares Nunes Monteiro, Markus Lange, Anderson Santos de Freitas, Luana do Nascimento Silva Barbosa, Johannes van Leeuwen, Jorge Emídio de Carvalho Soares, Rogério Eiji Hanada, Gerd Gleixner, Siu Mui Tsai

## Abstract

Soil phosphorus (P) is a limiting factor for vegetation growth in the Amazon rainforest, where plants depend on microorganisms for organic matter cycling and nutrient uptake. However, forest-to-agriculture conversion changes plant-microbe-soil interactions, affecting P cycling, which may additionally changed by land-use intensity. This study examined the 30-year effects of converting a primary forest into two contrasting systems: a low-intensity agroforest and a high-intensity citrus plantation. We investigated how microbial and water-extractable organic matter (WEOM) composition interacted with soil physicochemical attributes and P fractions (labile, moderately labile, non-labile, and residual). Agroforest soils retained physicochemical and enzymatic attributes similar to the primary forest, while soils of the citrus plantation showed increased P in all fractions due to fertilization and reduced soil organic matter content, mainly in deeper layers. Microbial and WEOM composition patterns reflected land-use, with agroforest representing an intermediate state between primary forest and citrus plantation. Proteobacteria and nutrient-rich WEOM were more abundant in the agroforest, whereas Ascomycota and nutrient-poor WEOM predominated the citrus plantation. Genes related to “P acquisition” were more abundant in agroforest soils, while genes related to “P-compound synthesis” were more abundant in citrus plantation. Labile P was negatively correlated with genes related to microbial metabolism, suggesting that reduced P availability may induce a boost in microbial activity for internal P-cycling. These findings demonstrate that forest-to-agriculture conversion strongly affects microbial functions, with responses aligning with land-use intensity and WEOM resource availability. Nonetheless, microbes adapt by shifting strategies: prioritizing mineralization and solubilization or favoring biosynthesis depending on P availability.

## 1. Introduction

The Amazon rainforest has experienced extensive deforestation in recent decades, disrupting essential plant-soil interactions that sustain ecosystem functions [1, 2]. The primary driver of land-use change in the region is the conversion of primary forests into agricultural and pastoral systems, often through slash-and-burn practices [3]. This transition alters the soil chemical, physical, and biological attributes and increases soil degradation [4, 5]. The combined loss of vegetation and soil organic matter, along with the nutrient volatilization, immobilization, and leaching, severely diminishes long-term soil health in these low-fertility soils [6]. Further, Amazonian soils are acidic and highly weathered, conditions that are often accompanied with the formation of stable P minerals through strong geochemical binding to iron and aluminum hydroxides, causing low concentrations of dissolved P [7]. As a result, P availability to plants is reduced, P becomes more recalcitrant, and microbial P cycling is altered [8, 9]. Given that most Amazonian soils are naturally nutrient-poor, P availability is considered a major constraint on plant productivity and ecosystem recovery following disturbance [10].

Therefore, sustainable agriculture is essential for preserving and maintaining ecosystem functions in the events of land-use change, including biogeochemical cycles. For instance, Amazonian agroforestry systems are often established on nutrient-poor soils, forcing plants to rely on strategies to overcome P limitations by recycling P through plant litter decomposition and microbial necromass [11]. In this sense, agroforestry integrates high plant diversity to optimize space and resource use, mimicking plant succession dynamics and above and belowground interactions as primary forests [12], which contrasts plantations. However, plant P uptake is limited to soluble inorganic P in form of orthophosphate, whereas most soil P exists in organic pools, such as litter, soil organic matter, and microbial biomass, as well as in inorganic forms adsorbed on soil particles with varying solubility [13]. However, despite their importance, the interactions between plants, microbes, and organic matter in P cycling remain poorly understood in the Amazon land use change systems.

In this context, soil microbiota and WEOM compounds are important indicators of shifts in biogeochemical processes. WEOM consists of a complex mixture of plant- and microorganism-derived metabolites containing CHNOPS elements (carbon, hydrogen, oxygen, nitrogen, phosphorus, and sulfur) [14, 15]. It functions as both a source and a sink for organic compounds, and depending on its composition and stoichiometry between elements, provides energy and nutrients for microbial activity [16]. Nutrient-rich WEOM typically presents low C:N ratio and high diversity of compounds, serving as a reservoir for microbially synthesized products [17]. In contrast, nutrient-poor WEOM is mainly composed of plant-derived compounds containing only carbon, hydrogen and oxygen [18]. Consequently, WEOM may play an essential role in the P cycle, making it fundamental to comprehend its interactions with soil microbial communities in systems undergoing land-use change in the Amazon. For example, plant-microbe interactions can contribute to nutrient retention and facilitate transformation of insoluble P fractions into labile forms available for plant uptake [9, 18]. As a result, WEOM stimulates microbial communities to perform nutrient cycling processes [19].

Microbial P-cycling involves hundreds of genes categorized into major “extracellular” and “intracellular” metabolic pathways [20]. The former is grouped into genes related to “P acquisition” which includes solubilization, mineralization, and transport of P. The later pathway includes genes related to synthesis of new “P-containing compounds”, such as purine and pyrimidine metabolism and oxidative phosphorylation [21, 22]. While these pathways are regulated by environmental conditions, the mechanisms governing microbial P acquisition and turnover in tropical soils remain poorly understood. Here, we investigate how land-use change, and management intensity affect microbial and WEOM composition, focusing on the potential expression of P-cycling genes involved in both extracellular and intracellular processes, and their relationship with different soil P fractions. We hypothesize that: (i) microbial and WEOM composition are strongly related to land use; and (ii) genes associated with both “P acquisition” and “P-compounds synthesis” pathways are influenced by microbial and WEOM composition and directly affects soil P fractions.

## 2. Material and Methods

### 2.1. Study site, Experimental Design and Management Intensity

The study was conducted in Manacapuru, Amazonas, Brazil (03°16’50” S; 60°30’17” W) (Fig. S1). The region has a Rainy Tropical (Amw) climate, according to the Köppen classification, with an annual average temperature of 28°C, humidity of 75-85%, and annual rainfall of 1,750–2,500 mm [23]. The soils are classified as Ferralsol [24]. The study sites are part of the “Ramal do Laranjal” land reform settlement, established in 1989 by the municipality of Manacapuru.

The agroforest system was established during the rainy season of 1992/1993 and 1993/1994 as a sustainable agriculture initiative by the National Institute of Amazonian Research (INPA). The site is managed by a local farmer and consists of three vertical strata: (i) *Theobroma grandiflorum* and *Paullinia cupana* in the lower stratum; (ii) *Euterpe precatoria* and *Bactris gasipaes* in the middle stratum; and (iii) *Bertholletia excelsa* in the upper stratum. The citrus plantation was established in the rainy season of 1997 using seedlings grafted with orange (*Citrus × sinensis*) as the graft, and mandarin lime (*Citrus × limonia*) as rootstock. Initial fertilization included 300 g of lime and 100 g of P_2_O_5_ per pothole, with mostly annual applications of 50 g each of CO(NH_2_)_2_, P_2_O_5_, and K_2_O per tree. A full description of the site’s management history is presented in the supplementary material (Table S1). The primary forest, used as a reference, is a highly preserved forest adjacent to the land reform settlement and has remained undisturbed for over 50 years.

Soil sampling occurred in May 2022 at the end of the wet season. A 60 × 60 m plot was established in each land-use system with five sampling points (four corners and center) to minimize edge effects. Five composite samples were collected per system at three depths (0–10 cm, 10–20 cm, and 20–30 cm) using a Dutch auger. Undisturbed samples for bulk density and porosity were collected with metallic volumetric rings (8 × 5 cm) using an Uhland-type auger. For molecular analysis, topsoil (0–10 cm) was collected in sterile centrifuge tubes, as this layer is a microbial activity hotspot [25]. Samples for chemical and physical analyses were stored at 4°C, while molecular analyses samples were preserved at -20°C.

### 2.2. Soil Chemical and Physical Analysis

Soil chemical attributes were analyzed following the procedures described [26]. Briefly, soil pH was measured in a CaCl_2_ solution (0.01 mol L^−1^). Soil organic carbon was quantified via oxidation using K_2_Cr_2_O_7_ solution (0.07 mol L^−1^). K^+^, Ca^2+^, and Mg^2+^ were extracted using ion exchange resins; K^+^ was quantified using a colorimetric method, while Ca^2+^ and Mg^2+^ were measured by atomic absorption spectrophotometry (PerkinElmer 3100, USA). SO_4_^2-^ was extracted with a Ca_3_(PO_4_)_2_ solution (0.01 mol L^−1^) and quantified by turbidimetry. Al^3+^ was extracted with a KCl solution (1.0 mol L^−1^) and quantified by titration with NaOH solution (0.025 mol L^−1^). Micronutrients, including Fe, Cu, Mn, and Zn, were extracted using diethylenetriaminepentaacetic acid (DTPA) and quantified via atomic absorption spectrophotometry. Cation exchange capacity (CEC) was estimated based on the sum of Ca^2+^, Mg^2+^, K^+^, Al^3+^, and H^+^.

Soil physical attributes were analyzed according to the methods described [27]. Shortly, soil bulk density was determined from undisturbed samples collected using metallic volumetric rings (8 × 5 cm). Soil macroporosity was measured through capillary rise water saturation by weighing samples and transferring them to a tension table at −6 kPa until hydraulic equilibrium was achieved. Microporosity was determined by drying the same samples at 105°C for 48 hours and reweighing them.

Soil enzymatic activity was assessed by quantifying the amount of ρ-nitrophenol released during the incubation of substrates analogs in 1 g of soil at 37°C for 1 hour, using a 96-well microplate reader (LMR-96 Loccus, São Paulo, Brazil). Enzymatic activity was measured using substrates analogs: β-glucosidase activity was measured at 410 nm using ρ-nitrophenyl β-glucopyranoside (PNG) [28]. Acid phosphatase activity was measured at 420 nm using ρ-nitrophenyl disodium phosphate (PNP) [29]. Arylsulfatase activity was measured at 410 nm using ρ-nitrophenyl sulfate (PNS) [30].

### 2.3. Phosphorus Fractionation Techniques

Soil phosphorus (P) fractionation was conducted following [31], with modifications [32]. Sequential extraction was performed using 0.5 g of air-dried soil in the following order: labile P, which is readily available for plant uptake, was extracted using 10 mL of Mehlich-3 solution, followed by shaking for 1 hour on an end-over-end tube rotator at 30 rpm, centrifugation at 3600 × g for 15 min, and collection of the supernatant. Moderately labile inorganic P was extracted with 10 mL of NaOH solution (0.5 mol L^-1^), followed by shaking for 1 hour, centrifugation for 15 minutes, and collection of the supernatant. Moderately labile organic P was obtained through autoclave digestion with 1 mL of H_2_SO_4_ (98%, analytical grade) and 0.75 g of (NH_4_)_2_S_2_O_8_ for 120 min. Non-labile P, primarily bound to calcium minerals, was extracted with 10 mL of HCl solution (1 mol L^-1^), followed by shaking and centrifugation. Additionally, total P content was determined using 0.1 g of soil. Samples were treated with 2 mL of H_2_SO_4_ (98%) and 2 mL of H_2_O_2_ (37%, analytical grade) and digested in heating blocks for 2 hours at 350°C. The P content in each fraction was quantified using the colorimetric method [33]. Residual P, consisting of highly recalcitrant inorganic and organic forms that are largely unavailable to plants, was estimated as the difference between total P and the sum of the extracted fractions.

### 2.4. Water-Extractable Organic Matter Extraction, and High-Resolution Mass Spectrometry

Water-extractable organic matter (WEOM) was obtained from topsoil (0–10 cm) following [34] with slight modifications. Briefly, 4.0 g of field-moist soil was mixed with 20 mL of KCl (2.0 mol L^−1^) (1:5 w/v), shaken at 20°C for 1 hour at 200 rpm, centrifuged at 8000 × g for 10 min, and filtered through a 0.45 µm glass fiber syringe filter (Whatman, Maidstone, UK). Extracts were acidified to pH 2.0 with HCl (37%, Sigma-Aldrich, USA), and the concentration of dissolved organic carbon (DOC) was quantified using a TOC analyzer (Shimadzu, Japan). The extract was cleaned using Agilent Bond Elute PPL Solid Phase Extraction cartridges (100 mg) [35]. Soil solution volume was adjusted based on DOC concentration to ensure 200 μg of organic carbon was loaded per cartridge. Compounds were eluted with ultrapure methanol, and SPE extraction efficiency was assessed on a Vario TOC cube analyzer (Elementar, Germany) [18].

High-resolution mass spectrometry (HR-MS) was performed using an Orbitrap Elite (Thermo Scientific), following [15, 36, 37]. The instrument was operated with direct injection of 100 µL of PPL-extract into the autosampler at a flow rate of 20 µL min^−1^, using a solvent mixture of ultrapure water and methanol (1:1). Electrospray ionization (ESI) was performed in negative mode with a voltage of 2.65 kV, without a separation column. Spectra were collected over a mass-to-charge (m/z) range of 100– 1000. Blank samples were included in all steps to identify and exclude background signals unrelated to WEOM.

An average of 100 scans per sample was combined into a single average mass spectrum and converted into spectral peaks using Thermo Xcalibur software (v. 3.0.63). ThermoRawFileParser software (v.1.7.4) [38] converted raw files to *mzML* format, retaining only m/z values with a signal-to-noise ratio >6 for further analysis.

### 2.5. Water-Extractable Organic Matter Compounds Analysis

Formula assignment was conducted using internal recalibration of spectral peaks with the *MFAssignR* package [39]. Ions with m/z values between 150–500 were considered. The sum formulae assignment was performed under the following conditions: ^12^C: 1–70, ^1^H: 2–160, ^16^O: 0–70, ^14^N: 0–7, ^32^S: 0–3, ^31^P: 0–3, and a tolerance of ± 0.5 ppm. This process resulted in the assignment of 9710 formulae across 15 samples. From these formulae, we determined key molecular metrics, including the hydrogen-to-carbon ratio (H/C), oxygen-to-carbon ratio (O/C), carbon-to-nitrogen ratio (C/N), modified aromaticity index (AImod), aromatics metabolites [40], nominal oxidation state of carbon (NOSC) and the difference between double bond equivalency and oxygen atom count (DBE-O) [41].

### 2.6. DNA Extraction and Sequencing Protocols

Total DNA was extracted from topsoil (0–10 cm) using 0.25 g of soil. Extraction was performed with PowerSoil Pro Kit (Qiagen, Hilden, Germany), following manufacturer’s protocol. DNA quantity and quality were measured using a Nanodrop 2000c spectrophotometer (Thermo Fisher, USA), and DNA integrity was confirmed by 1% agarose gel electrophoresis at 90 V for 30 min.

For amplicon sequencing, 16S rRNA gene and ITS region were targeted to analyze prokaryotic and fungal communities, respectively. The V3–V4 hypervariable region of the 16S rRNA was amplified using the 515F/806R primers [42], while the ITS region was amplified using the ITS1f/ITS2 primers [43]. Sequencing was carried out on the Illumina NextSeq platform (Illumina, San Diego, USA).

Shotgun metagenomics was performed on the Illumina HiSeq 2500 platform (Illumina, San Diego, USA) with the HiSeq Reagent Kit v.4, following the manufacturer’s recommendations. DNA libraries were constructed using 2 µg of DNA per sample, fragmented to sizes lower than 500 bp. DNA fragments were amplified by PCR using P5 and P7 primers for multiplexing and subsequently purified and quantified using a Qubit 2.0 fluorometer (Invitrogen, Carlsbad, USA). Finally, libraries were demultiplexed and sequenced with paired-end reads of 2 × 100 bp.

### 2.7. Microbial Community and Functioning Analysis

Processing of amplicon sequencing data, including quality control, reconstruction of 16S rRNA and ITS genes, and taxonomic annotations, was conducted using the *nf-core/ampliseq* pipeline (v.2.11.0) [44, 45]. The pipeline was run with default settings, employing Cutadapt (v.1.16) to remove primers, QIIME2 for sequence import, and the DADA2 algorithm [46] for sample processing. Sequences with a quality score below 30 and PCR chimeras were discarded, and a minimum read frequency of three was applied to exclude singletons and doubletons. Taxonomic annotation was performed using the SILVA database (v.138) for prokaryotes and the UNITE database (v.9.0.) [47] for fungi. Taxa classified as Archaea and Eukaryote were removed from the prokaryotic dataset. The sequences are available at the NCBI Sequence Read Archive under the identification PRJNA1244318.

Metagenomics functional profile was assessed with the DiTing pipeline (v.0.9) [48] to analyze the relative abundance and occurrence of metabolic and biogeochemical pathways related to microbial P cycling. Briefly, reads were assembled using MEGAHIT (v.1.1.3) [49], and open reading frames (ORFs) were predicted from contigs using Prodigal (v.2.6.3) [50]. ORFs annotations were conducted using the KEGG ortholog database with HMMER3 (v.3.4). The relative abundance of KOs was obtained by mapping the ORF against the metagenomic reads, with results expressed as transcripts per million (TPM).

### 2.8. Statistical Analysis and Data Interpretation

All statistical analyses and visualizations were conducted in R (v.4.4.1). Soil attributes were tested for normality and homoscedasticity of residuals to assess their suitability for parametric analysis. When assumptions were met, differences between land-use systems were tested using one-way ANOVA, followed by Tukey’s HSD test (p < 0.05) for post-hoc comparisons. Otherwise, the Kruskal-Wallis test was applied, followed by Dunn’s post-hoc test (p < 0.05).

Principal Component Analysis (PCA) was performed using the *factoextra* package [51] to evaluate differences in soil chemical attributes across land-use systems. Principal Coordinate Analysis (PCoA) based on Bray-Curtis dissimilarity was performed to examine microbial and WEOM composition, using the *vegan* package [52]. Differences in microbial and WEOM compositions were assessed with PERMANOVA using the *pairwiseAdonis* function (p < 0.001) and interpreted by the coefficients of determination (R^2^). The relative abundance of bacteria, fungi, and WEOM were transformed in centered log-ratio (*clr*) and means were compared by the Kruskal-Wallis test, using the *ALDEx2* package [53]. Heatmaps based on Spearman correlation (p < 0.0001) were generated to evaluate the relationships between P-cycling genes and P fractions, and between WEOM and P fractions, using the *corrplot* package.

Partial Least Squares Path Modeling (PLS-PM) [54] was performed to explore the effects of microbial and WEOM composition on P-cycling pathways and their relationship with P fractions. A complete description of the paths and hypothesis behind the model is described in the supplementary files (Table S2). The *plspm* package [55] was used to estimate the strength of path coefficients (i.e., direct effects) and coefficients of determination (R^2^) for linear relationships among variables, using 1000 bootstraps for validation. Model assumptions were checked via *unidimensionality* (Cronbach’s alpha > 0.7) using the *outerplot* function, the commonality of latent variables (commonality > 0.49) through cross-loading scores. The final model structure was visualized using the *innerplot* function.

## 3. Results

### 3.1. Land-use change impacted soil attributes and phosphorus fractions

Principal component analysis (PCA) revealed distinct soil chemical differences across the land-use systems, with the first two principal components explaining 66.6% of total variation (Fig. 1A). The first axis separated the citrus plantation from the primary forest and the agroforest, mainly due to higher concentrations of magnesium (Mg) and calcium (Ca) in the first soil depth (Fig. 1B). Citrus soils showed higher pH, as well as elevated concentration of manganese (Mn) and zinc (Zn), while the primary forest and agroforest soils showed higher concentrations of aluminum (Al^3+^), boron (B), and cation exchange capacity (CEC) (Fig. 1B; Table S3). The second axis mainly showed the effect of soil depth, with upper layer having higher concentrations of soil organic matter (OM), iron (Fe), potassium (K) (Fig. 1B; Table S3).

**Fig. 1.**
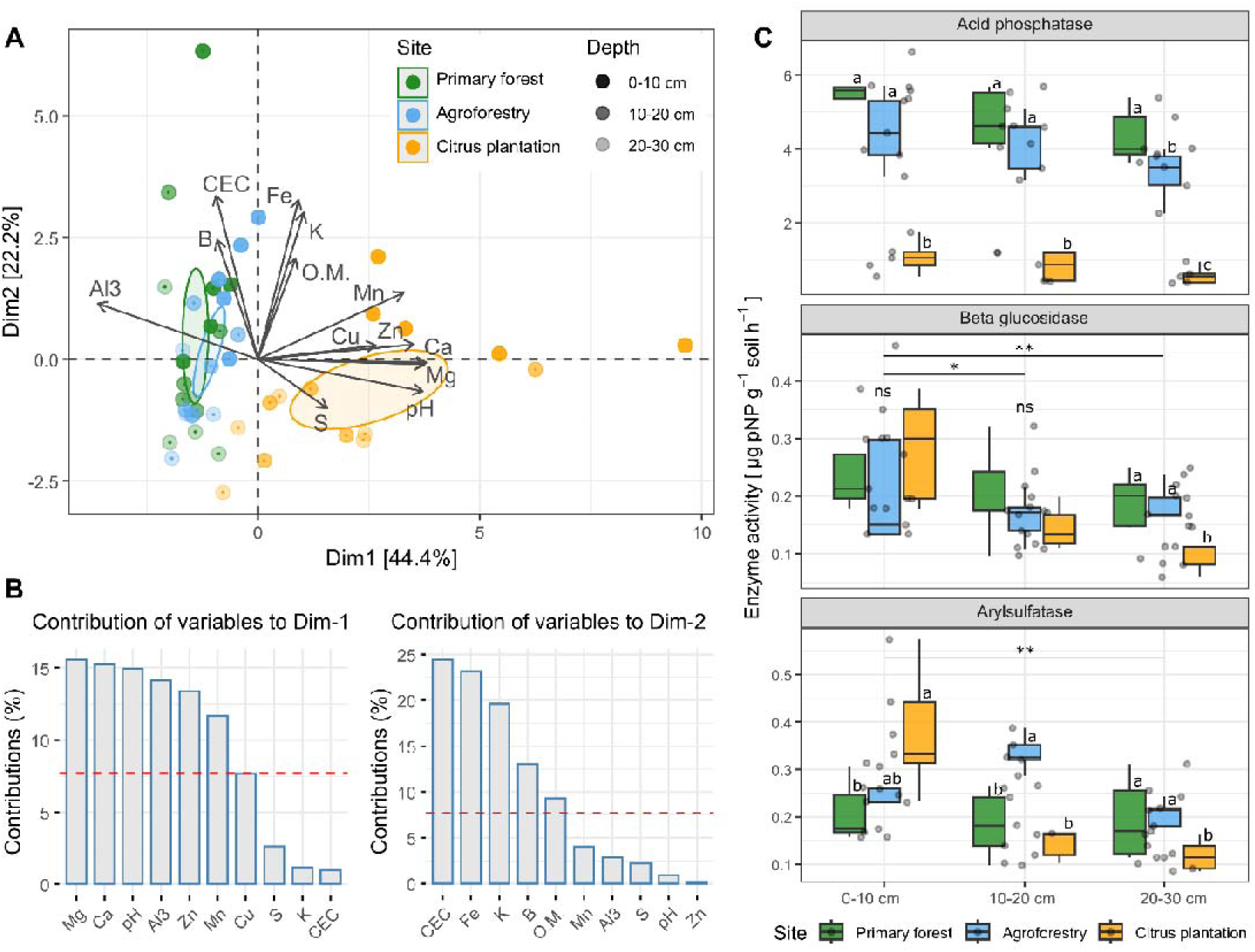
(A) Principal component analysis (PCA) of soil chemical attributes across land-use systems. Ellipses represent the 95% confidence interval. (B) Variables contribution to the first and second dimension of the PCA. Dashed red line indicates the significant threshold of variables. (C) Soil enzymatic activity across land-use systems at different soil depths. Boxplots show mean, median, and 95% confidence interval (*n* = 5). Letters indicate significant differences between land-use systems at the same soil depth by the Kruskal-Wallis test followed by Dunn’s post-hoc (p < 0.05). Asterisks indicate significant differences between soil depths by one-way ANOVA (** = p < 0.01; * = p < 0.05).

Soil biological attributes also varied across land-use systems and soil depths (Fig. 1C). Acid phosphatase activity was highest in forest soils, followed by the agroforest and citrus plantation at all depths. Beta-glucosidase activity showed no significant differences among sites in the top two soil layers (0–10 cm, 10–20 cm) but was significantly higher in primary forest and agroforest soils compared to citrus plantation at 20–30 cm. In contrast, arylsulfatase activity was highest in citrus plantation at 0–10 cm but lower than forest and agroforest at deeper layers. Soil physical attributes differed mainly in deeper layers (Fig. S2). Total porosity was significantly higher in agroforest soils than in forest and citrus plantation soils in the second and third depths, while soil bulk density was significantly higher in citrus at 10– 20 cm and 20–30 cm. No significant differences were observed in the topsoil.

Phosphorus fractions varied significantly after long-term forest conversion (Fig. 2). Citrus plantation soils exhibited higher total P content and higher concentration of P in all extractable fractions compared to primary forest and agroforest in all three layers. In the topsoil layer (0–10 cm), the agroforest had a similar total P content to citrus plantation but showed the lowest concentration in the deepest layer (20–30 cm). The majority of P across all sites was found in the organic fraction at all soil depths. Citrus plantation soils also contained higher levels of inorganic moderate labile and labile phosphorus at all depths. In contrast, primary forest and agroforestry soils had higher residual phosphorus content compared to citrus plantation.

**Fig. 2.**
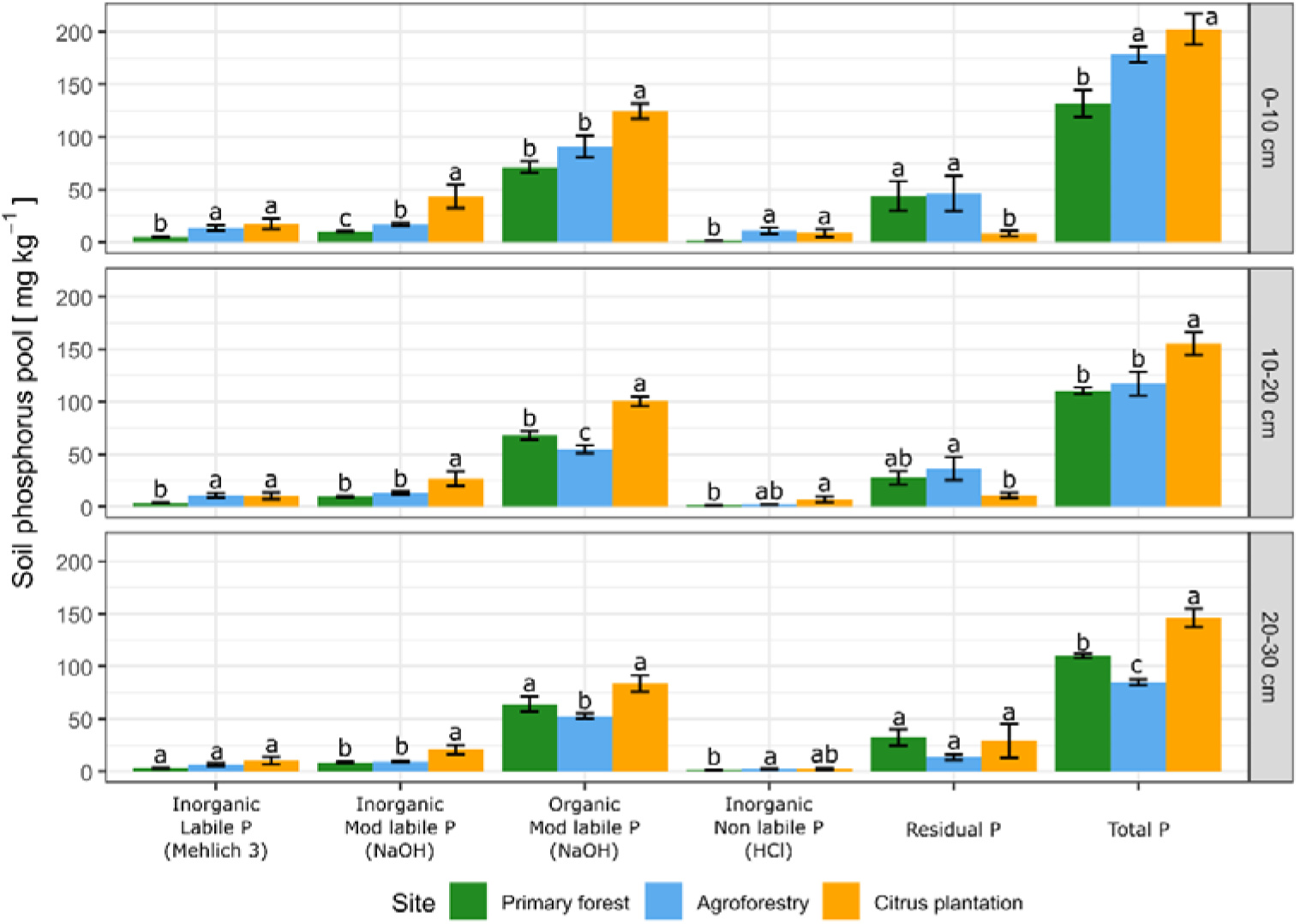
Soil phosphorus fractions in different soil depths across land-use systems. Fractions are shown in their sequential extraction order. Data are mean ± SE (*n* = 5). Letters indicate significant differences between land-use systems by the Kruskal-Wallis test followed by Dunn’s post-hoc (p < 0.05).

### 3.2. Microbial and WEOM composition across different land-use systems

A total of 3966 bacterial ASVs were identified, with an average of 82098 reads per sample. Proteobacteria was significantly more abundant in primary forest compared to other sites, Acidobacteria was more abundant in agroforest, although differences were not significant, while Chloroflexi was significantly higher in citrus plantation (Fig. S3). For fungi, 7724 ASVs were identified, averaging 111620 reads per sample. Ascomycota and Basidiomycota comprised over 70% of the fungal community, though many taxa remained unclassified at phylum level. Ascomycota was the most abundant phylum for all sites although differences were not significant, while Basidiomycota was significantly more abundant in primary forest and agroforest, and Glomeromycota was higher in citrus plantation (Fig. S3).

Microbial composition differed significantly across land-use systems after long term forest-to-agriculture conversion (Fig. 3). Bacterial composition showed the most pronounced differences between primary forest and citrus plantation (PERMANOVA R^2^ = 0.45), followed by primary forest and agroforest (R^2^ = 0.41), and agroforest and citrus plantation (R^2^ = 0.33) (Fig. 3A). Bray-Curtis distance analysis revealed that bacterial composition was more homogeneous in agroforest compared to citrus plantation due to lower dissimilarity indices (Fig. S4). Similarly, fungal communities exhibited the most pronounced differences between primary forest and citrus plantation (R^2^ = 0.32), while the differences between primary forest and agroforest, as well as between agroforest and citrus plantation, were identical (R^2^ = 0.24) (Fig. 3B). However, the Bray-Curtis distance revealed that the fungal composition was more heterogeneous in agroforest soils than in citrus plantation (Fig S4).

**Fig. 3.**
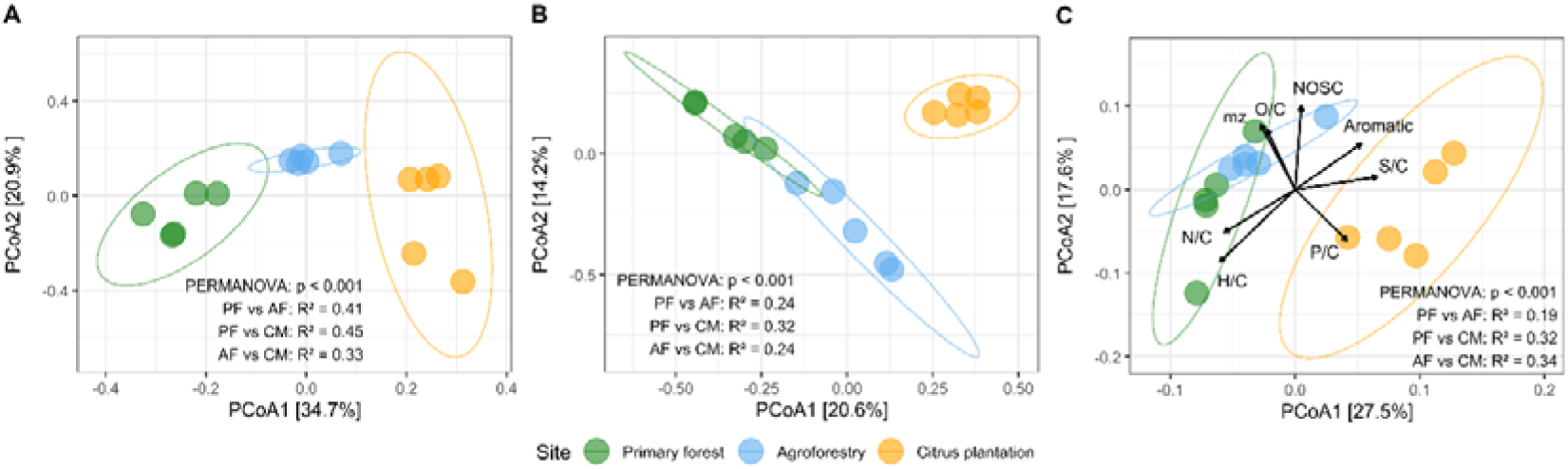
(A) Bacterial, (B) fungal community, and (C) Water-extractable organic matter (WEOM) composition across land-use systems represented by a Principal Coordinate Analysis (PCoA) based on Bray-Curtis dissimilarity. Arrows indicate intensity-weighted WEOM parameters, fitted as vectors. Coefficients of determination (R^2^) denote the proportion of variance explained by the model.

For WEOM, 9710 molecular formulas were identified, comprised primarily by CHO (carbon, hydrogen, and oxygen), and CHNO (carbon, hydrogen, oxygen, and nitrogen) formulas, which together represented 96% of the total assigned formulas in the WEOM composition (Fig. S3). Although there were no significant site differences, CHO formulas increased by an average of 12.4%, and CHNO formulas decreased by an average of 13.3% in citrus soils compared to primary forest soils. WEOM composition closely reflected microbial community patterns (Fig. 3C). The strongest differences in WEOM composition occurred between agroforest and citrus plantation (R^2^ = 0.34), followed by comparisons between the primary forest and citrus plantation (R^2^ = 0.32), and between the primary forest and agroforest (R^2^ = 0.19). Among the WEOM parameters, the hydrogen-to-carbon (H/C) ratio had a higher relationship with primary forest soils, while phosphorus-to-carbon (P/C) ratio was more related to citrus plantation soils. The mass-to-charge ratio (m/z) and the nominal oxidation state of carbon (NOSC) were by tendency separated both agroforest and primary forest soils from the citrus plantation.

### 3.3. Phosphorus cycling genes and pathways across different land-use systems

A total of 92 genes involved in P-cycle were identified across land-use systems (Fig. 4A). The most abundant genes were *pyrG* (encoding the orotidine-5′-decarboxylase), *pstS* (phosphate-binding protein), *guaA* (glutamine amidotransferase), and *prsA* (post-translocation molecular chaperone). Land-use influenced the majority of P-cycling genes, with distinct patterns observed: genes upregulated in primary forest were downregulated in citrus plantation, while genes downregulated in primary forest were upregulated in citrus. Agroforest exhibited intermediate levels of gene expression compared to the other two systems.

**Fig. 4.**
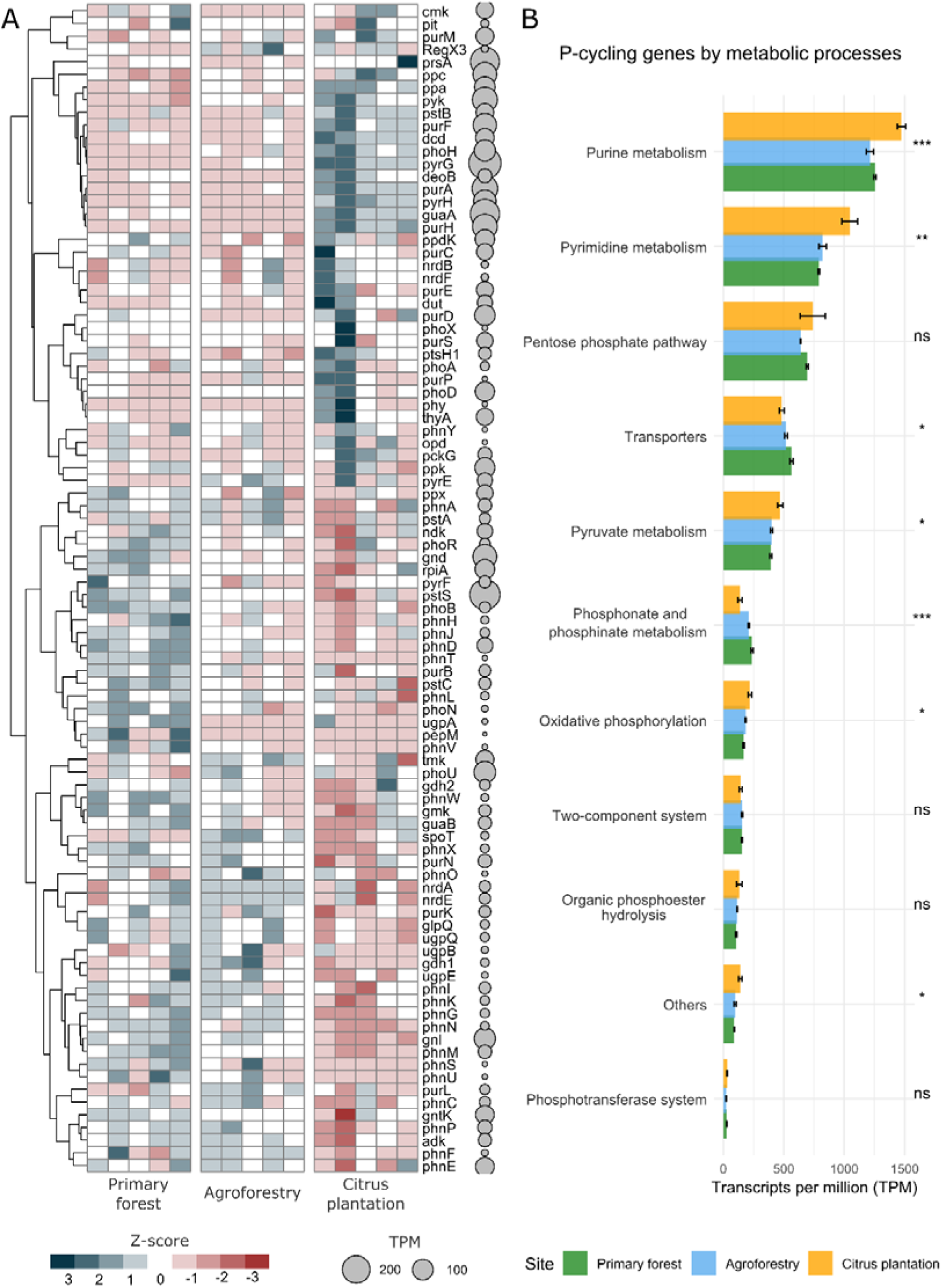
(A) Heatmap showing the abundance of phosphorus cycling genes in different land-use change systems. Colors from blue to red represents a range from high to low abundance of transcripts per million (TPM) normalized by z-score. (B) Barplot showing the metabolic pathways involved in microbial phosphorus cycle represented by TPM. Data are mean ± SE (*n* = 5). Asterisk indicate significant differences among land-use systems by one-way ANOVA (*** = p < 0.001; ** = p < 0.01; * = p < 0.05).

The highest transcript abundance was associated with purine metabolism, followed by pyrimidine metabolism, the pentose phosphate pathway, and transporters (Fig. 4B). Genes related to the “P-compounds synthesis” pathway, such as the purine and pyrimidine metabolism, and the oxidative phosphorylation were significantly upregulated in citrus plantation compared to primary forest and agroforest. In contrast, genes related to the “P acquisition” pathway which included transporters, phosphonate and phosphinate metabolism were significantly higher in primary forest than in agroforest and citrus plantation.

### 3.4. Environmental drivers of phosphorus cycling genes

P-cycling pathways and WEOM composition had significant relationships with different soil P fractions (Fig. S5). The organic and residual P fractions showed contrasting correlation patterns. The organic P fraction was positively correlated with several P-cycling pathways, such as pyrimidine metabolism, purine metabolism, and oxidative phosphorylation, as well as with WEOM composition such as CHP and CHOSP compounds. In contrast, the residual P fraction showed positive correlations with pathways including the pentose phosphate pathway, transporters, and phosphonate and phosphinate metabolism. Further correlations showed that P-cycling pathways were also affected by the WEOM composition (Fig. S6). The strongest positive correlations were observed between pyrimidine metabolism and CHP compounds (ρ = 0.71), followed by two-component system and CHNP (ρ = 0.65), and purine metabolism and CHP (ρ = 0.60). Conversely, the strongest negative correlations were observed between phosphonate and phosphinate metabolism with CHP (ρ = -0.75), followed by organic phosphoester hydrolysis and CHNOS (ρ = -0.63).

PLS-PM analysis revealed that both microbial and WEOM composition influenced the abundance of genes associated with the “P acquisition” and “P-compound synthesis” pathways, which in turn affected soil P fractions (Fig. 5A and Table S4). Path analysis showed that microbial composition had a stronger overall effect on P-cycling genes than WEOM composition. Specifically, the first PCoA axis for bacteria and fungi, representing microbial composition, was positively correlated with the “P-compounds synthesis” pathway (0.58) and negatively correlated with the “P acquisition” pathway (-0.76). In contrast, WEOM composition showed weaker correlations with these pathways (0.57 and -0.17, respectively). Microbial composition also had a stronger positive correlation with the labile P fraction (1.44) compared to the moderate labile P fraction (0.83), while WEOM composition did not significantly influence either P fraction. The two microbial P-cycling pathways showed contrasting effects on P availability. The “P acquisition” pathway was positively correlated with moderately labile P (0.60), whereas the “P-compound synthesis” pathway was negatively correlated with labile P (-0.78).

**Fig. 5.**
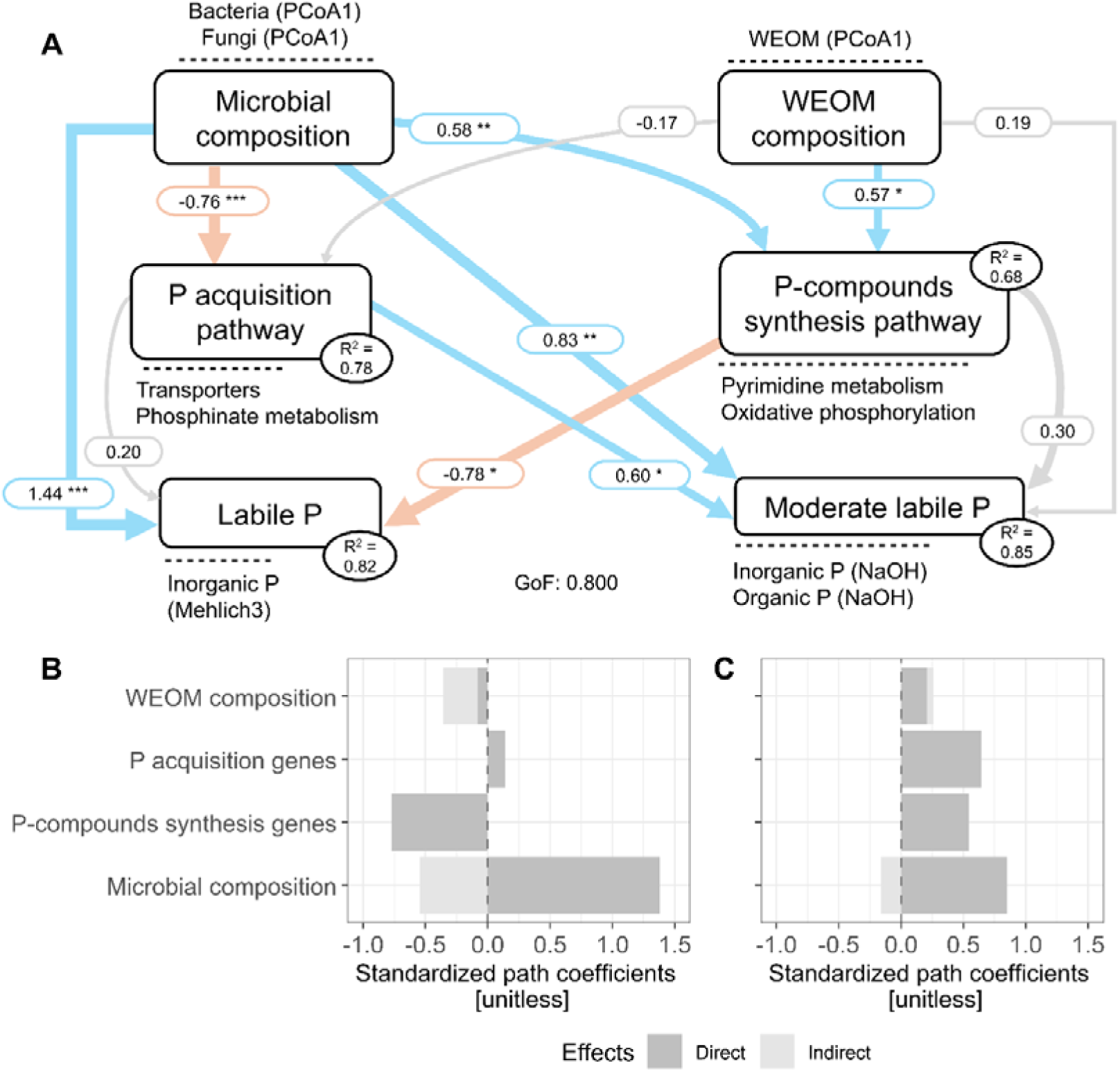
(A) Partial least squares path modeling (PLS-PM) showing the direct effects of microbial and water-extractable organic matter (WEOM) composition on P-cycling pathways and P fractions. Direct and indirect effects of the environmental variables on (B) moderate labile P and (C) labile P. Boxes represent latent variables used in the PLS-PM and variables near the dashed lines represents the observed variables. R^2^ values are the coefficients of determination of the endogenous latent variables. Blue and red arrows indicate significant positive and negative effects, respectively. Grey arrows indicate non-significant effects. Arrow width indicates the relationship strength. Numbers near the arrows indicate significant standardized path coefficient. Asterisks indicate significant effects (* = p < 0.05; ** = p < 0.01; *** = p < 0.001).

## 4. Discussion

This study investigated the influence of microbial and WEOM composition on P-cycling genes and P fractions across different Amazonian land-use systems. As hypothesized, land-use change significantly altered soil physicochemical attributes, microbial composition and function, and the WEOM profile, primarily driven by changes in nutrient availability. However, the conversion of forest to agroforest mitigated the impacts on both biotic and abiotic soil attributes compared to the more intensive citrus plantation. P-cycling genes showed distinct correlations with soil P fractions: the “P acquisition” pathway positively correlated with moderate labile P, while the P-compounds synthesis” pathway negatively correlated with labile P. These results highlight the adaptive role of soil microbes to regulate P cycling processes in response to changes in WEOM composition and resource availability (Roth et al., 2019; Lange et al., 2024). Overall, our findings suggest that land management practices can influence microbial function and improve P-use efficiency, even in nutrient-poor soils like those in the Amazon.

### 4.1. Microbial and organic matter composition patterns after long-term land-use change

Land-use change has well-documented impacts on Amazonian soil attributes and microbial composition [9, 56, 57]. However, this study provides a novel perspective on ecosystem functioning by demonstrating that WEOM composition closely reflects microbial composition providing important insights into biogeochemical processes. For instance, the conversion of primary forest into agroforest maintained a homogeneous bacterial community with a lower dissimilarity index, while converting into citrus plantation resulted in the homogenization of fungal communities (Fig. S4). Likewise, WEOM composition became more homogeneous in agroforest compared to primary forest and citrus plantation, suggesting that a considerable portion of organic compounds may originate from bacterial activity. These compositional shifts are likely influenced by reduced soil organic matter content [58] but also by high management intensity practices, such as fertilization and liming in citrus plantation [59, 60]. Moreover, microbial divergence aligned with environmental drivers, as bacterial communities tends to respond to nutrient availability, and fungal communities responds to aboveground vegetation [12]. These results emphasize the potential to influence the soil microbiome through vegetation manipulation, as promoting plant diversification enhances a broader fungal community, and improving nutrient availability supports bacterial community development [61].

In this context, the observed microbial patterns reinforce the hypothesis that microbial and organic matter composition are tightly linked in response to land-use change. For example, the increased abundance of Chloroflexi is often correlated with reduced organic carbon and nitrogen levels [62], while the higher abundance of Acidobacteria corresponds to more acidic pH and organic matter-rich conditions [63, 64]. This group is especially important in highly weathered soils due to their tolerance to aluminum toxicity [65], a common condition observed in tropical soils. Such shifts in microbial distribution may indicate declining soil health, as these changes involves transitions from oligotrophic to copiotrophic taxa – e.g., from Ascomycota to Basidiomycota, or from Acidobacteria and Chloroflexi to Proteobacteria and Actinobacteria [66, 67]. Although microbial life strategies may not fit accurately into a binary classification of copiotrophic versus oligotrophic, especially at the phylum level [68], our data suggest that forest-to-agriculture conversion tends to favor taxa adapted from high-to low-carbon soils. This shift is likely driven by reductions in plant diversity and soil carbon, which can also decrease nutrient use efficiency [69]. However, monocultures, such as citrus plantation, can amplify these effects.

These patterns were also reflected in the WEOM composition, supporting the idea that organic compound diversity is shaped by specific plant and microbial inputs, as well as soil attributes [70, 71]. Our results suggest that WEOM transformation is primarily driven by the bacterial community [70, 72], as nutrient-rich compounds (e.g., CHNP formulas) were more prevalent in the agroforest, while nutrient-poor compounds (e.g., CHO formulas) dominated in citrus plantation (Fig. S3). These compositional changes were evident in H/C, N/C ratios and NOSC in agroforest, which is often indicated as a result of greater microbial processing [73]. In summary, land-use change alters both microbial and WEOM composition, transitioning from nutrient-rich, microbially derived compounds to nutrient-poor, plant-derived compounds. However, transition to agroforestry appears to mitigate the adverse effects of land-use change, maintaining functionality close to that of primary forests.

### 4.2. Phosphorus cycling genes and phosphorus fractions after long-term land-use change

The long-term conversion of forest to agriculture altered the soil P fractions through changes in physical, chemical, and biological soil attributes. Differences in land management intensity, such as increased plant diversity in agroforestry or the use of fertilizers in citrus plantation further expanded these changes. Consequently, soil microbial communities and their associated P-cycling genes also diverged, reinforcing differences in soil P fractions.

However, despite annual fertilization in citrus plantation, total P concentrations remained critically low (< 200 mg kg^-1^), which is concerning for agriculture in the Amazon, where over 60% of soils are classified as low-P Ferralsols [11, 74]. In the agroforest, total P in topsoil was similar to citrus plantation but significantly lower in deeper layers, suggesting more efficient nutrient mining in the subsoil layers. However, soil P accumulated primarily in topsoil of agroforest, suggesting that these systems are highly dependent on soil health to ensure effective nutrient cycling [75, 76]. Moreover, agroforest soils showed higher litter biomass, litter P content (Fig. S7 and S8), and phosphatase activity than citrus plantation, indicating enhanced P release through organic matter decomposition and stimulation of P-cycling microbes [77]. This was supported by a greater abundance of microbial genes involved in P mineralization and transport, reflecting efficient P recycling even under limited nutrient availability. The WEOM composition also aligned with the hypothesis linking resource availability in organic matter to microbial activity. Under nutrient-poor WEOM conditions, internal P-cycling appeared to occur mainly via microbial metabolism, while nutrient-rich WEOM stimulated microbial P-cycling processes through mineralization and transporters genes. As a consequence, the enrichment of P- and N-containing WEOM over CHO-WEOM in the agroforest is likely byproducts of microbial activity [18].

Nevertheless, sequential P fractionation provides a more accurate assessment of plant-available P than identifying which soil properties affects the P fractions or the chemical forms of different P pools [78]. This may explain the stronger correlation between microbial communities and labile P, as well as the negative correlation between labile P and nutrient-poor WEOM. These results indicate that under P-limited conditions, microbial communities can access non-labile and residual P fractions and convert them into more bioavailable forms for plants. Although sequential fractionation alone does not fully capture biogeochemical complexity of P cycling [79], integrating it with metagenomics and mass-spectrometry offers a more comprehensive understanding. Accordingly, our findings suggest that labile and moderately labile P fractions are important pools regulated by key P-cycling pathways, such as purine and pyrimidine metabolism, along with CHP and CHOSP compounds. These processes are strongly associated with copiotrophic taxa such as Actinobacteriota and Proteobacteria [80], which contribute to organic matter turnover and restoration of soil health. Given that soil organic matter is a key determinant of labile P availability [81, 82], we emphasize that both quantity and composition of organic matter determinates microbial access to resource for effective P cycling.

## 5. Conclusions

Long-term conversion of primary forest into agroforest and citrus plantation has significantly altered soil attributes and microbial functions associated to P cycling. Both microbial and WEOM compositions were strongly shaped by land use, which in turn influenced the potential for microbial P-cycling. Genes associated to “P acquisition” (e.g., transporters, phosphonate metabolism) were more abundant in agroforest soils compared to citrus plantation and had a positive correlation with moderate labile P fractions. In contrast, genes linked to “P-compounds synthesis” (e.g., pyrimidine metabolism, oxidative phosphorylation) were enriched in citrus plantation and negatively correlated with the labile P fraction. Despite the higher organic P fraction in citrus plantation soils, acid phosphatase activity was greater in the primary forest and the agroforest, suggesting that fertilization may suppress microbial P mineralization.

Agroforest with low management intensity consistently exhibited intermediate levels between the primary forest and citrus plantation with high management intensity, supporting both plant productivity and nutrient cycling in nutrient-poor soils. These patterns highlight the adaptability of microbial communities in accessing and using P, with functions prioritized based on resource availability. This was further evidenced in WEOM composition: primary forest and agroforest soils were enriched with nutrient-rich compounds, whereas citrus plantation soils contained more nutrient-poor compounds. By integrating genomic, metabolic, and environmental data, this study offers novel insights into the microbial mechanisms of P-cycling in tropical soils and highlights the role of WEOM in further supporting this process. Overall, managing microbial functions and WEOM composition offers a promising path for improving P-use efficiency and promoting sustainable agriculture in the Amazon, where P is a finite and limited resource. However, our findings represent a temporal snapshot and may vary with seasonality, soil types, and different land management.

## Acknowledgments

We would like to express our gratitude to Eduardo Vasconcelos for his maintenance of the agroforest and the citrus plantation site since the beginning of the experiment, to Lucas Franhani Alexandre for his help with the soil sampling campaign, and to Vitor Fernando Barro for his help with the soil enzyme analysis. The authors would also like to express their gratitude to Fundação de Amparo à Pesquisa do Estado de São Paulo (FAPESP) and Fundação de Amparo à Pesquisa do Estado do Amazonas (FAPEAM) for their financial support for the project (2020/08927-0). GLM is thankful to FAPESP for the PhD scholarship (2022/05561-0) and the international scholarship (2024/06334-2). GGTNM thanks FAPESP for his scholarship (2024/02443-1). ML gratefully acknowledges funding by the Zwillenberg-Tietz Foundation.

## Author Contributions

**GLM**: conceptualization, formal analysis, investigation, methodology, visualization, writing – original draft, writing – review & editing; **GGTNM**: formal analysis, validation, writing – review & editing; **ML**: validation, writing – review & editing; **ASF**: validation, writing – review & editing; **LNSB**: investigation, validation; **JvL**: data curation, writing – review & editing; **JECS**: data curation; **REH**: funding acquisition, writing – review & editing; **GG**: supervision, methodology, validation, writing – review & editing; **SMT**: conceptualization, funding acquisition, project administration, resources, supervision, validation, writing – review & editing.

## Data availability statement

The datasets generated during and/or analyzed during the current study are available in the Zenodo repository: https://doi.org/10.5281/zenodo.15147497.

